# A fast and inexpensive plate-based NGS library preparation method for insect genomics

**DOI:** 10.1101/2023.11.24.568434

**Authors:** Lauren Cobb, Erik de Muinck, Spyros Kollias, Morten Skage, Gregor D. Gilfillan, Markus A. K. Sydenham, Shuo-Wang Qiao, Bastiaan Star

## Abstract

Entomological sampling and storage conditions often prioritise efficiency, practicality and conservation of morphological characteristics, and may therefore be suboptimal for DNA preservation. This practice can impact downstream molecular applications, such as the generation of high-throughput genomic libraries, which often requires substantial DNA input amounts. Here, we investigate a fast and economical Tn5 transposase tagmentation-based library preparation method optimised for 96-well plates and low yield DNA extracts from insect legs stored under different conditions. Using a standardised input of 6ng DNA, library preparation costs were significantly reduced through the 6-fold dilution of a commercially available tagmentation enzyme. Costs were further suppressed by direct post-amplification pooling, skipping quality assessment of individual libraries. We find that reduced DNA yields associated with ethanol-based storage do not impede overall sequencing success. Furthermore, we find that the efficiency of tagmentation-based library preparation can be improved by thorough post-amplification bead clean-up which selects against both short and large DNA fragments. By lowering data generation costs, broadening the scope of whole genome studies to include low yield DNA extracts and increasing throughput, we expect this protocol to be of significant value for a range of applications in the field of insect genomics.

## Introduction

In response to widespread declines in insect diversity (reviewed in Wagner et al. 2021), and the subsequent severe implications for ecosystem functioning (reviewed in van der Sluijs 2020), large-scale international insect survey and sampling schemes have been put in place (e.g. Potts 2020). While the primary interest of such schemes is to monitor abundance and distribution changes, the application of genomic approaches to insect specimens can also be used to investigate spatial drivers of population trends, habitat connectivity and local adaptation (reviewed in Saunders et al. 2020). These approaches rely on high-throughput sequencing (HTS) methods, which generally have considerably greater DNA quality requirements compared to more traditional PCR-based methods. However, entomological samples are often collected and stored using a variety of techniques which form a trade-off between practicality, morphological requirements and molecular demands. While samples should ideally be stored immediately following tissue death under optimal conditions (Graham et al. 2015), this is not always feasible during extended field periods. For instance, entomological specimens are often temporarily stored in warm fieldwork vehicles until sampling has concluded, and those captured by high-efficiency trapping methods, such as pitfall and flight interception traps, can remain in traps on site for several days, or even weeks, prior to collection and processing (Knuff et al. 2019). In addition, as entomological specimens are important biological resources with myriad applications in the fields of taxonomy and ecology, standard collection and storage methods are designed to prioritise morphological preservation. Given these considerations, alcohols such as ethanol are commonly used as both killing agents and preserving fluids (e.g. Potts 2020). While this approach helps to prevent physical deterioration, it does not fully inhibit DNA degradation due to relatively high water content (Lindahl 1993, Soniat et al. 2021). The use of highly concentrated ethanol (>96%) can improve DNA preservation, although ethanol concentration may decrease due to evaporative loss. In addition, entomological specimens preserved in high concentration ethanol can become overly brittle, impacting morphological analyses (King and Porter 2004). Ethanol-based sampling and storage nevertheless remains the cheapest and most practical method compared to other options such as flash freezing with liquid nitrogen, and is therefore widespread (e.g. Frampton et al. 2008, Moreau et al. 2013, Sousa et al. 2023). Library preparation methods which are compatible with material collected and stored using suboptimal preservation protocols are therefore preferred when working with entomological specimens.

An additional obstacle to the use of entomological specimens for HTS is the destructive nature of DNA extraction: certain protocols require the sample be physically ground or beaten with beads, while most call for tissue digestion through the use of corrosive chemicals and enzymes (e.g. Qiagen 2020). Some ‘non-destructive’ insect DNA extraction methods have been proposed, whereby the entire specimen is incubated during the digestion step and subsequently retained, allowing for the preservation of the whole insect (Thomsen et al. 2009, Castalanelli et al. 2010, Tin et al. 2014, Santos et al. 2018). However, this practice may result in some morphological damage such as pigment loss (Tin et al. 2014, Korlević et al. 2021), and its impacts on long-term preservation (i.e. > 10 years) remain unclear. Importantly, while re-extractions of the same biological material may produce viable genomic DNA extracts, the first extraction is often the most effective in terms of DNA yield and endogenous DNA content (Cavill et al. 2022). Using the entire specimen therefore limits any future DNA applications, reduces the long-term biological value of entomological collections, and fails to incorporate ethical guidelines for the destructive sampling of biological specimens (Pálsdóttir et al. 2019). To most effectively preserve insect specimens, for both future entomological applications and future DNA extractions, a more conservative solution would be to sacrifice a small piece of tissue such as a leg (Cavill et al. 2022). This approach leaves the majority of the insect untouched, although the low sample volume and consequent low DNA yield may constrain specific downstream molecular applications.

Although the cost of whole genome sequencing has decreased by several orders of magnitude since its inception (Shendure et al. 2017), library preparation often remains expensive and now represents a substantial proportion of the overall cost of whole genome data generation. For instance, methods whereby DNA is physically fragmented are widely adopted, yet can be expensive, time consuming, and often require high DNA input (Tvedte et al. 2021). Alternatively, tagmentation-based library preparation has streamlined the process by combining DNA fragmentation and adapter ligation into a single reaction facilitated by Tn5 transposase, and typically calls for lower DNA input (Adey et al. 2010, Jones et al. 2023). Commercial Tn5 transposase kits, however, remain relatively expensive, and their dependence on the use of undisclosed reagents inhibits methodological optimisation. Several adaptations to the tagmentation method have been developed, such as the in-house production of Tn5 transposase (Picelli et al. 2014). Despite significantly reducing costs, in-house enzyme production can be both time and resource intensive, may not be always possible and therefore lacks broad-scale practicality. Further modifications include reduction of reaction volumes, replacement of reagents with cheaper alternatives and the incorporation of magnetic bead-linked transposomes, highlighting the potential of matching low DNA input amounts with diluted transposase concentrations to increase both throughput and affordability (Baym et al. 2015, Bruinsma et al. 2018, Jones et al. 2023). Here, we explore the use of a commercially available hyperactive Tn5 transposase (Diagenode) to create high-throughput sequencing libraries from low-yield DNA extracts obtained from entomological specimens.

Specifically, we investigate the effectiveness of an in-house developed, fast and cheap Tn5 tagmentation-based library preparation method, using insect DNA extracted by applying minimally destructive methods to samples collected and stored using two common preservation protocols. First, we assessed the impact of varying tagmentation reaction time and Tn5 transposase enzyme concentration on library fragment size distribution. We then rescaled our protocol to fit 96-well plates and a standardised DNA input of 6ng. Finally, we developed a double-sided post-amplification clean-up protocol in order to optimise library fragment size distribution and thus improve sequencing efficiency. We find that this economical, easily scalable library preparation protocol opens multiple opportunities for genomic studies, particularly in the cases of non-model organisms with limited funding, small organisms with low per-sample DNA yield or biological samples stored under non-optimal conditions for DNA preservation.

## Materials and Methods

### Specimens, sampling and storage conditions

122 samples of two different bumblebee species (*Bombus lapidarius* and *Bombus pascuorum*) were stored following one of two protocols (see Table 1). All specimens were sampled in 2017 and 2022 by immediate submersion in 96% ethanol. They were then stored and transported in fieldwork vehicles for several weeks during the summer, before being dried, pinned and stored at room temperature with no preservative (Dry Protocol, *n* = 82). Samples stored using the Ethanol Protocol (*n* = 40) were comprised of single legs which were removed from the specimens collected in 2017 prior to drying, and subsequently stored in 96% ethanol at −18°C.

**Table 1.** Bumblebee specimen sampling and storage features. Details of two bumblebee sample storage protocols, including year sampled, killing agent, storage preservative and storage temperature, and the number of legs used for DNA extraction. Also presented are the sample sizes from each group used in two different sets of analyses: DNA concentration comparison of DNA extracts, and library preparation and sequencing statistics.

| Protocol | Year sampled | Killing agent | Storage preservative | Temperature | Specimens (n) | One leg extracted (n) | Two legs extracted (n) | DNA concentration comparison (n) | Library preparation and sequencing (n) |
| --- | --- | --- | --- | --- | --- | --- | --- | --- | --- |
| Dry | 2017 / 2022 | Ethanol | None | Ambient | 82 | 21 | 61 | 21 | 82 |
| Ethanol | 2017 | Ethanol | Ethanol | -18°C | 40 | 40 | 0 | 40 | 8 |

### DNA extraction

DNA was extracted from either 1 or 2 mid or hind legs, including the coxa, femur, tibia, and tarsomeres, using the Qiagen DNeasy Blood & Tissue Kit. Samples from the Ethanol Protocol were washed with molecular-grade water in order to remove any ethanol residue. All samples were incubated in 380μL digestion buffer (300μL ATL buffer, 50μL DTT, 30μL proteinase K) at 56° C and 350 RPM overnight (minimum 17 hours). Following overnight digestion, 7μL RNase A and 300μL AL buffer were added to the lysate, followed by a room temperature incubation (15 minutes) and the addition of 360 μL 96% ethanol. DNA was subsequently bound and purified following the manufacturer’s recommendations, then eluted in AE buffer (105μL, 90μL or 80μL) and stored at −18°C. DNA concentration was then estimated using the Qubit dsDNA Assays (Thermofisher) and standardised to 1ng/μL.

Initially, DNA extractions of dry-stored specimens were performed on single legs (n=21). In order to consistently avoid very low DNA yields, two legs were used for the remaining 61 extractions, along with the addition of 5μL 1M CaCl_2_ solution to the digestion buffer to aid tissue digestion.

### Library preparation

Two 8-sample Illumina compatible trial libraries were generated to evaluate the impact of two variables on DNA fragment size distribution: 1) tagmentation reaction time (Trial 1) and 2) Tn5 transposase enzyme concentration (Trial 2). For Trial 1, two *B. pascuorum* DNA extracts were incubated for five, six, seven or eight minutes with 1μL of undiluted Loaded Tn5 Tagmentase (Diagenode, C01070012). For trial two, the same DNA extracts were incubated with diluted Tagmentase (either a 2X, 4X, 6X or 8X dilution) for 7 minutes.

Following these trials, the protocol was adapted for a 96-well plate setup, and 96 libraries were generated using a 7 minute incubation and 6X Loaded Tn5 Tagmentase dilution. The libraries were built using extracts with DNA concentrations of at least 1ng/μL, consisting of 82 dry-stored samples, 8 ethanol-stored samples, and 6 additional samples which were either found to be misidentified bumblebee species, or stored using a different protocol. Following PCR amplification, the libraries were pooled and cleaned using two alternative clean-up protocols (Pool SP and Pool S4) in order to optimise fragment size distribution for sequencing (see also Supplementary Material for full Tn5 tagmentation library preparation, PCR and post-amplification clean-up protocol.).

### Tn5 tagmentation library preparation and PCR

All DNA extracts were standardised to 1ng/μL. 6μL DNA extract were incubated (55°C) with 5μL of 2X Tagmentation Buffer (Diagenode, C01019043) and 1μL of Loaded Tn5 Tagmentase (Diagenode, C01070012, using various dilutions, see above). After 5, 6, 7 or 8 minutes (see above), 3μL 0.2% SDS was added to stop the tagmentation reaction, followed by a 5 minute room temperature incubation. Post-tagmentation PCR (12 cycles) was done in 40μL reactions using KAPA HiFi (Roche). The manufacturer’s recommendations were followed by adding 24μL Mastermix, 1μL N7 Primer (1μM) and 1μL N5 primer (1μM) to each 15μL reaction. After amplification, 6μL of each of the 96 wells was pooled (without quantification) before proceeding to clean-up (AMPure XP, Beckham Coulter).

### Post-amplification clean-up and sequencing

The Trial 1, Trial 2 and Pool SP libraries were cleaned once, removing small DNA fragments by using a 0.6 DNA/bead ratio, following the manufacturer’s recommendations. Pool S4 was built from the same PCR reaction as Pool SP, but cleaned twice, using different bead ratios to remove both large and small DNA fragments. Firstly, a 0.5 DNA bead/ratio was used to remove large DNA fragments. Secondly, a 0.65 DNA/bead ratio was used to remove smaller DNA fragments and shift the fragment size distribution to peak between 500 and 1000 bp. This “double sided” clean-up process was performed twice, and the final library was eluted in 40μL molecular grade water. Fragmentation distribution profiles were visualised using a Fragment Analyzer (Advanced Analytical). Pool SP was sequenced using an Illumina Novaseq 6000 SP flowcell, generating 923,253,536 read pairs passing filters and Pool S4 was sequenced using an Illumina Novaseq 6000 S4 flowcell and generated 2,799,713,996read pairs passing filters. Base calling and demultiplexing were performed with RTA v3.4.4 and bcl2fastq v2.20.0.422 in both cases.

Read data were aligned using PALEOMIX v1.2.14 (Schubert et al. 2014) to BomLapEIv1 (https://www.ncbi.nlm.nih.gov/datasets/genome/GCA_936014575.1/) and BomPasc1.1 (Crowley et al. 2023) after using AdapterRemoval/2.3.1 (Schubert et al. 2016) and the mem algorithm as implemented in BWA/0.7.17 (Li 2013). Only reads with a minimum MapQ value of 15 were retained and considered endogenous.

### Statistics

The DNA yield comparison between dry-stored and ethanol-stored specimens was carried out using a t-test. Mann-Whitney-Wilcoxon tests were used to compare sequencing statistics between a) dry-stored and ethanol-stored specimens and b) Pool SP and Pool S4, excluding the endogenous DNA comparison between Pool SP and Pool S4, for which a t-test was used. All statistical analyses were performed using base R 4.3.1.

## Results

### Sample storage impact on DNA yield

One leg was used to extract DNA from samples stored in ethanol at −18°C (*n*=40) and either one leg (*n*=21) or two legs (*n*=61) were used from samples stored dried at ambient temperatures (Table 1). When comparing DNA yields of extracts generated from comparable tissue volumes (a single leg), we found that significantly more DNA is obtained from the dry-stored specimens than the ethanol-stored specimens (Figure 1). Given that DNA yields from ethanol-stored specimens were low, we chose to prioritise dry-stored specimens for library preparation.

**Figure 1.**
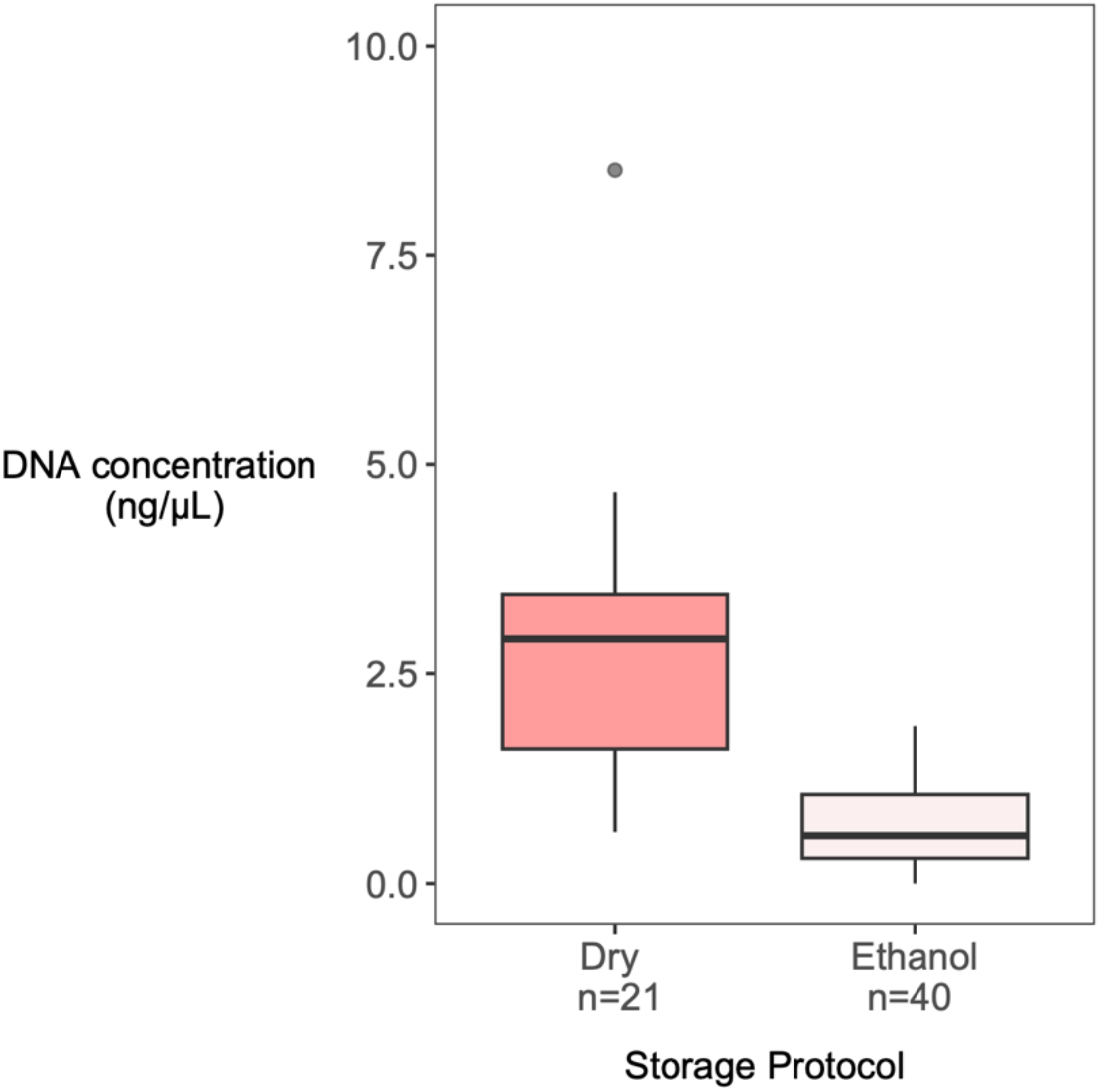
DNA extract concentration measurements from single bumblebee legs (*B. pascuorum* and *B. lapidarius*) stored using 2 different methods. Specimens were sampled in 2017 from various locations in Norway, Germany and Denmark and were either stored dry (at room temperature) or in ethanol (at −18°C). See Materials and Methods for details regarding sample collection and DNA extraction. Following extraction, DNA was eluted in 105μL. DNA yields from dry-stored specimens (mean = 2.84±1.72) were significantly greater than from those stored in ethanol (mean *=* 0.69±0.50; t = 5.46, df = 20.59, P < 0.001).

### Tn5 transposase library preparation optimisation

Changing tagmentation reaction time had minimal impact on fragment size distribution, with all libraries showing similar distribution profiles and fragment size peaking slightly above 300 bp (see Figure 2a). Fragmentation size distribution profiles of libraries built using 2X, 4X and 6X transposase enzyme dilutions were broader, and showed a modest increase in the proportion of long DNA fragments with each ascending dilution. This increase becomes more pronounced with the use of an 8X dilution (see Figure 2b). We therefore generated libraries in a 96-well plate using a 7 minute incubation time and 6X Loaded Tn5 Tagmentase dilution. These libraries were then amplified (12 cycles), after which 5μL from each library was pooled to perform bead clean-up on all libraries combined using two different approaches.

**Figure 2.**
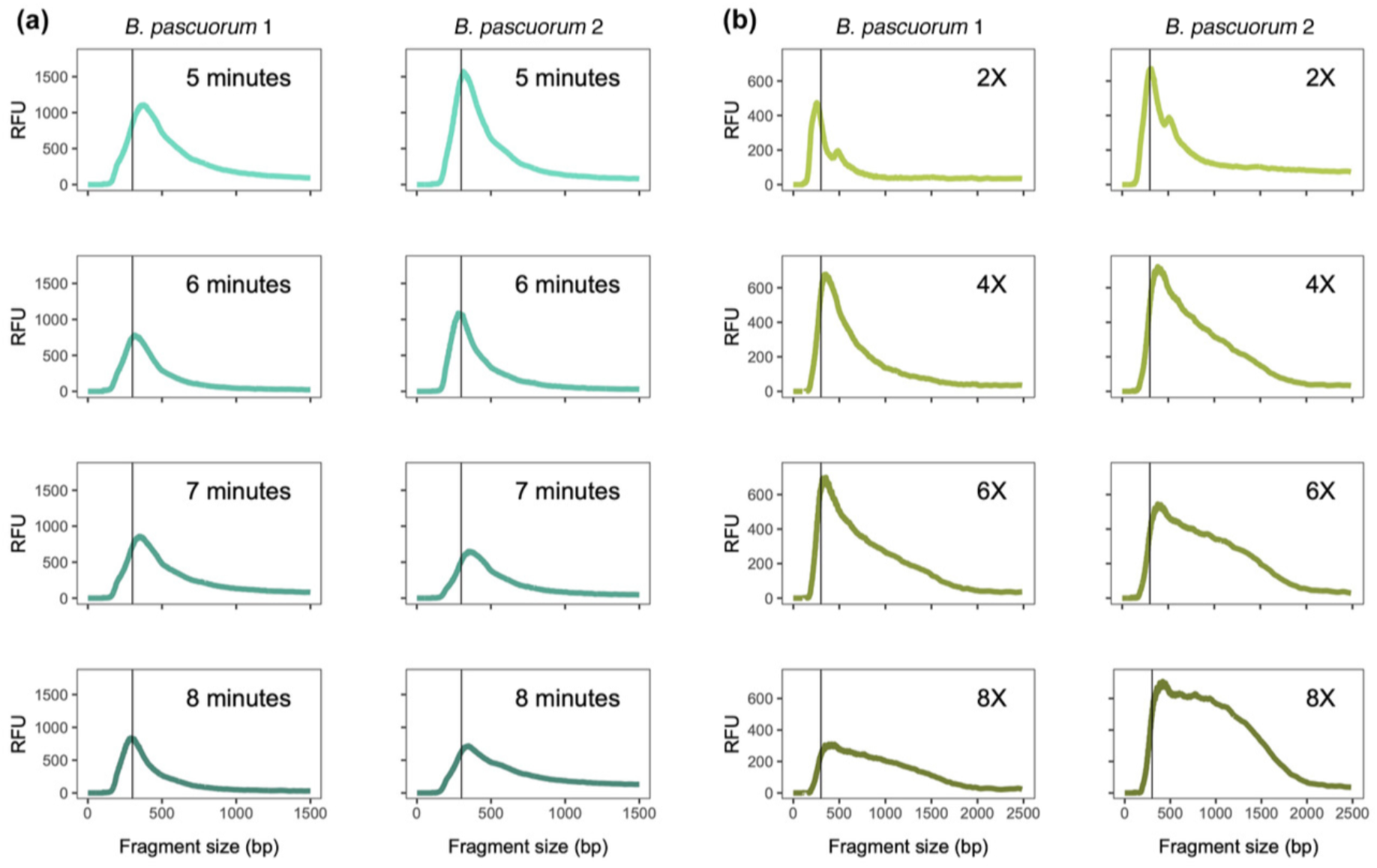
Fragment size distribution profiles of DNA libraries built from a standardised input of 6ng DNA from each of two *B. pascuorum* specimens, using different tagmentation reaction times and transposase dilutions. **(a)** Libraries were built using four different tagmentation reaction times (5 minutes; 6 minutes; 7 minutes; 8 minutes) with a 1X transposase dilution. **(b)** Libraries were built using four different transposase dilutions (2X; 4X; 6X; 8X) with a 7 minute tagmentation reaction. RFU (Relative Fluorescence Units) signifies relative amount of DNA. A vertical line indicating a fragment size of 300 bp is included to assist visual comparison of the different treatments. See Materials and Methods for details regarding library preparation protocol.

### Post-amplification clean-up optimisation

Following initial bead clean-up, we obtained a fragment size distribution situated between 400 bp and 1500 bp (Pool SP, Figure 3), a much broader range than the tight distribution around 600 bp recommended for Illumina DNA sequencing (Illumina DNA Prep Reference Guide, 2020). Pool SP was sequenced on an Illumina Novaseq SP flowcell with paired-end 150 bp reads, with a large number of reads collapsing into a single read (see further below). We therefore performed a more stringent clean-up on the original pooled libraries to tighten fragment size distribution, removing both very short and very long DNA fragments, resulting in a narrow distribution peaking around 600 bp (Pool S4, Figure 3). Pool S4 was subsequently sequenced on one quarter of an Illumina 6000 S4 flowcell, also with paired-end 150 bp reads.

**Figure 3.**
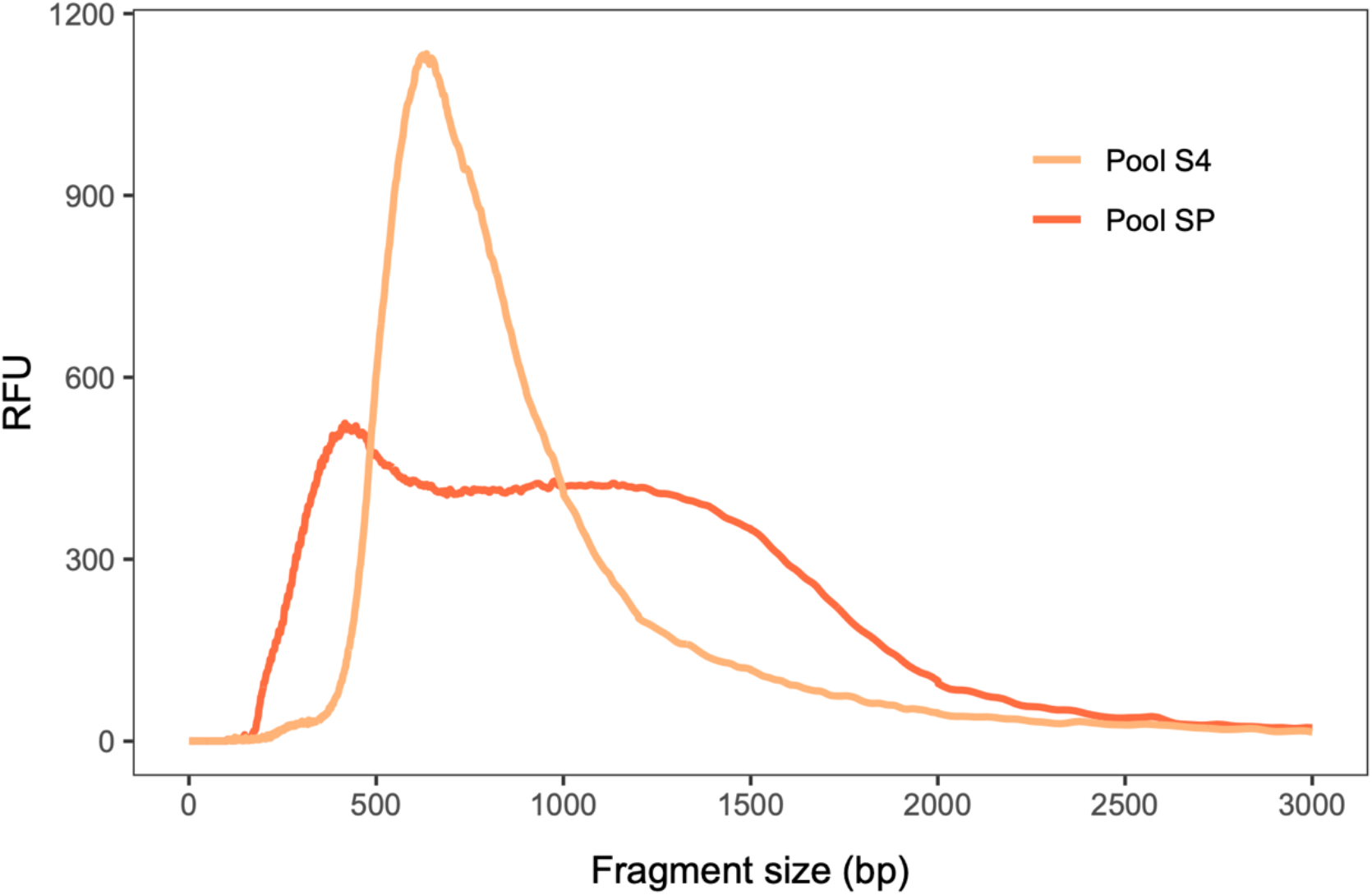
Comparison of fragment size distribution of 96 pooled bumblebee DNA libraries which were cleaned up with different AMPure XP protocols: a single clean-up with a 0.6 DNA/bead ratio (Pool SP), and a double-sided clean-up using two ratios (0.5 DNA/bead ratio followed by 0.65 DNA/bead ratio) which was performed twice (Pool S4). RFU (Relative Fluorescence Units) signifies relative amount of DNA.

### Sequencing results

We obtained 795,694,049 reads ranging between 261,576 and 23,087,451 per specimen for Pool SP, while Pool S4 produced significantly more reads: 2,409,119,984, ranging between 835,001 and 96,527,020 per specimen (W = 429, P = <0.001; Table 2; Figure 4a). Furthermore, Pool SP contained a significantly higher number of collapsed reads (W = 8100, P = <0.001; Figure 4b), slightly lower endogenous DNA proportion (t = −25.89, df = 89, P = <0.001; Figure 4c), and slightly higher overall read length (W = 8012, P = <0.001; Figure 4d) compared to Pool S4. Overall, Pool S4 resulted in higher coverage than Pool SP (W = 365, P = <0.001; Figure 4e), which was predominantly driven by the higher sequencing yield (Figure 4a). Nonetheless, when calculating the ratio of coverage against sequencing yield (calculated as coverage divided by number of paired reads), the coverage achieved by Pool S4 was on average greater than that of Pool SP by a factor of 1.45±0.12. This increase in efficiency for Pool S4 can be explained by both its reduced proportion of collapsed reads, leading to the mapping of more nucleotides per pair, and its higher proportion of endogenous DNA, leading to more unique mapped sequences per pair.

**Figure 4.**
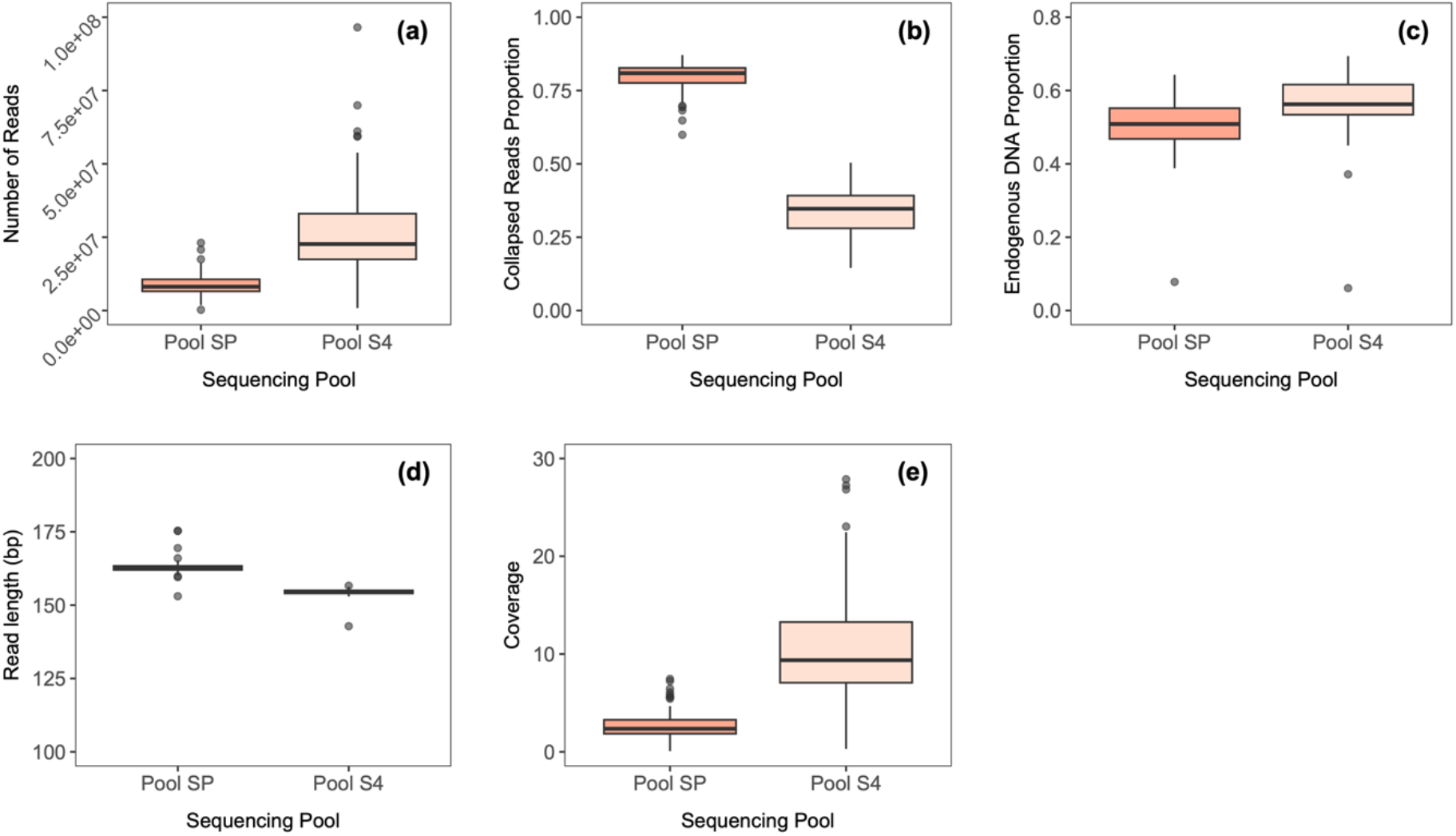
Sequencing comparisons of two pools of 90 bumblebee libraries. Bumblebee specimens were sampled between 2017 and 2022 from various locations in Norway, Germany, Denmark and Sweden. Both pools were generated from the same parallel library session. Pool SP was cleaned up with a 0.6 DNA/bead ratio and was sequenced on an Illumina Novaseq SP flowcell. Pool S4 was cleaned up using a double-sided size selection protocol, with a 0.5 DNA/bead ratio clean-up followed by a 0.65 DNA/bead ratio clean-up (twice) and was sequenced on ¼ Illumina Novaseq S4 flowcell. The boxplots compare **(a)** number of reads; **(b)** proportion of collapsed reads; **(c)** endogenous DNA proportion (unique reads only); **(d)** read length and **(e)** coverage.

**Table 2.**
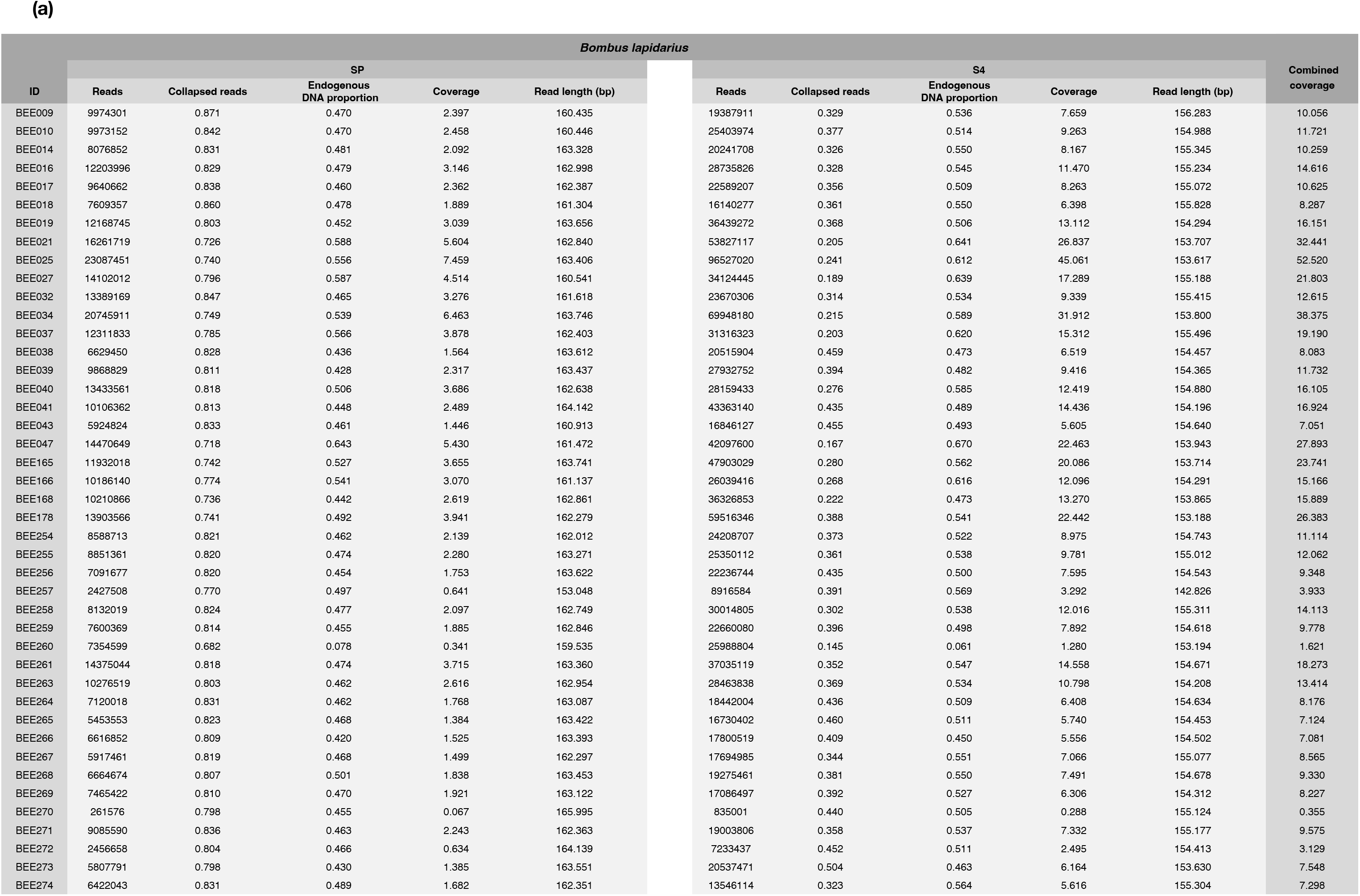
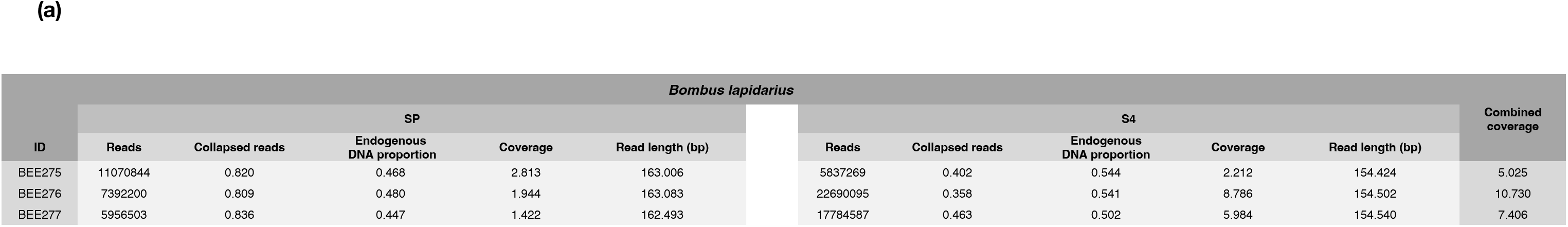
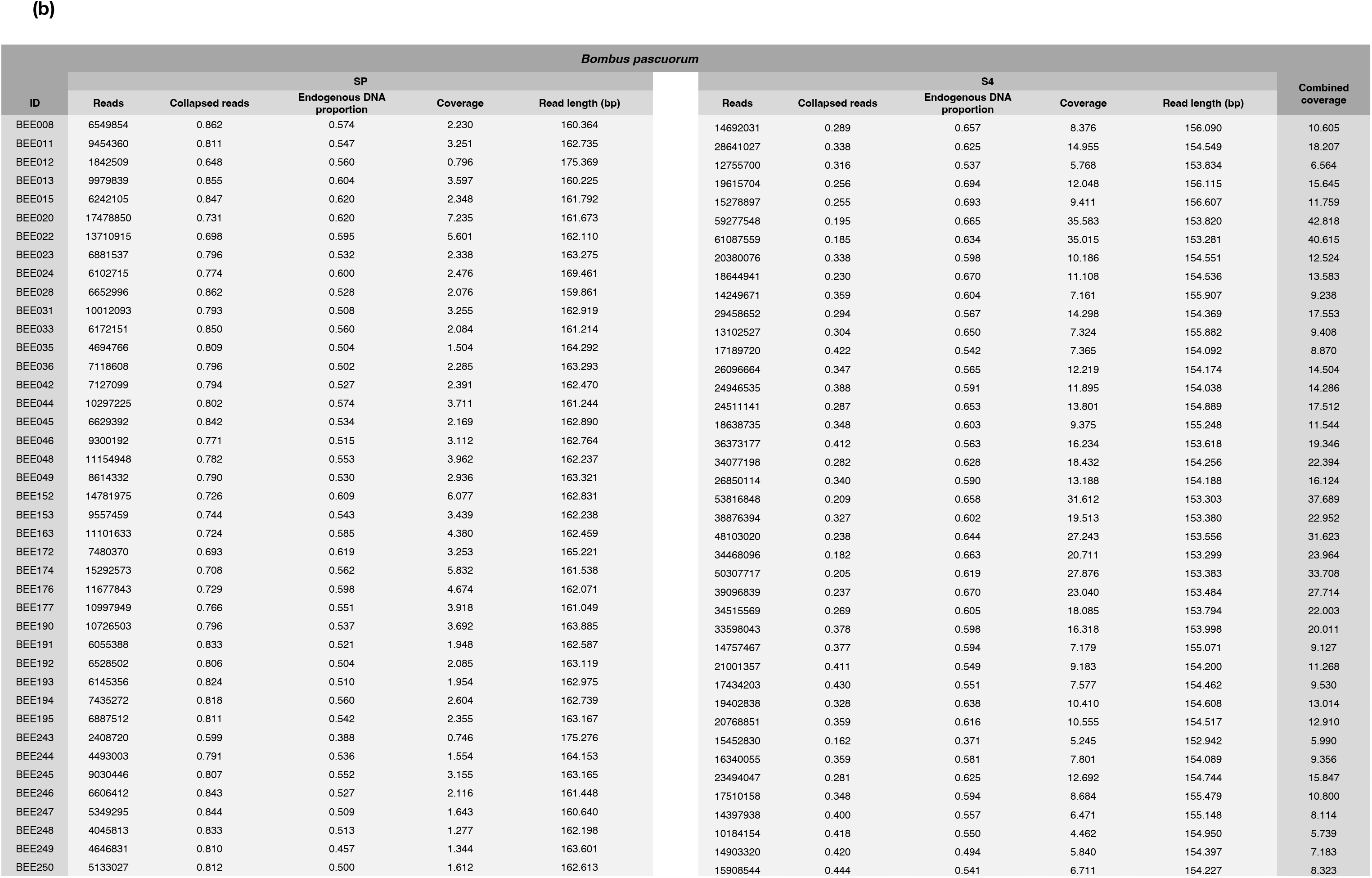
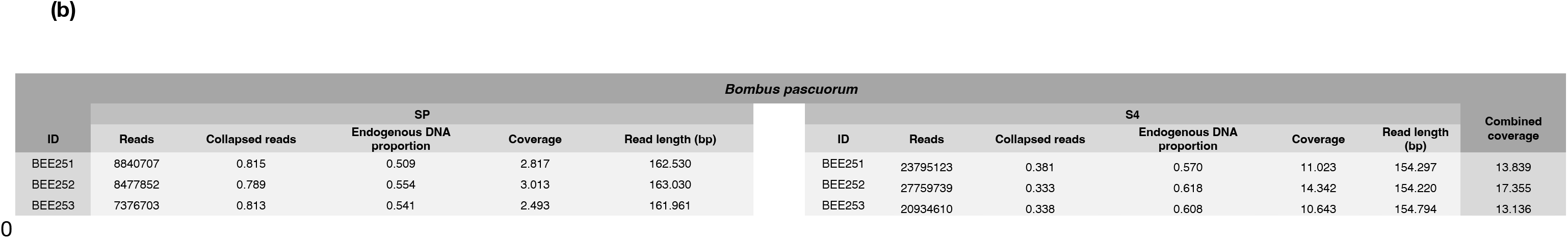
a & b. Sequencing statistics for two sequencing pools (SP and S4) of 90 specimens of two bumblebee species: (a) *Bombus lapidarius* (*n* = 46) and (b) *Bombus pascuorum* (*n* = 44). Bumblebee specimens were sampled between 2017 and 2022 from various locations in Norway, Germany, Denmark and Sweden. Libraries for both pools were generated simultaneously. The SP pool was cleaned up with a 0.6 DNA/bead ratio and was sequenced on an Illumina Novaseq SP flowcell. The S4 pool was cleaned up using a double-sided size selection protocol, with a 0.5 DNA/bead ratio clean-up followed by a 0.65 DNA/bead ratio clean-up (twice) and was sequenced on ¼ Illumina Novaseq S4 flowcell. Statistics presented for each sequencing run are specimen ID, number of reads, proportion of collapsed reads, clonality, endogenous DNA proportion, coverage, and read length (bp). Also presented is combined per-specimen coverage of both sequencing runs.

Ethanol-stored samples consistently produced preferable sequencing statistics, with significantly more paired reads (W = 553, P = <0.001; Figure 5a; see Supplementary Material), fewer collapsed reads (W = 1850, P = 0.007; Figure 5b), higher endogenous DNA proportions (W = 402, P = <0.001; Figure 5c), and similar read length (Figure 5d), leading to significantly higher overall coverage (W = 598, P = <0.001; Figure 5e).

**Figure 5.**
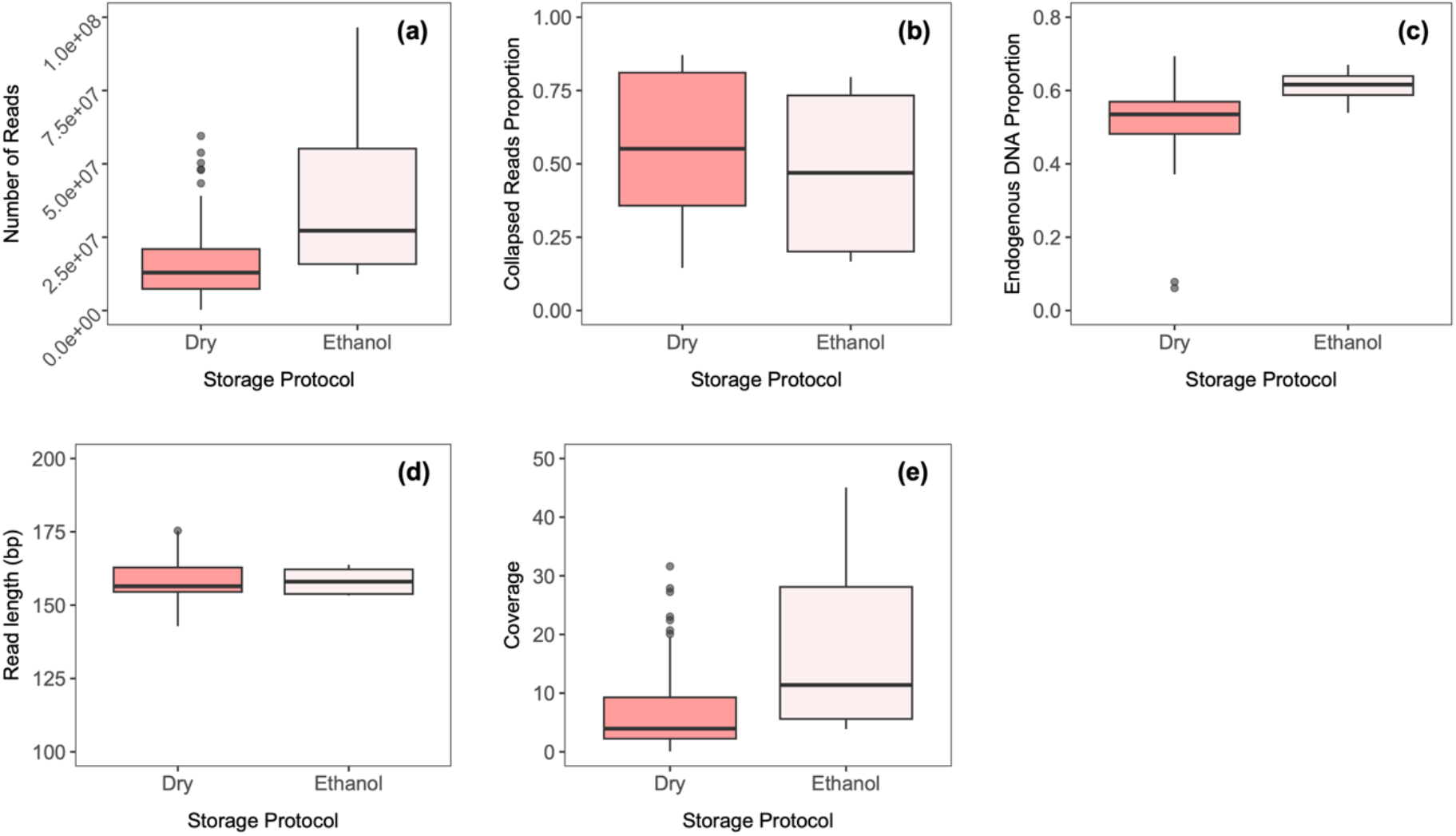
Sequencing comparisons of two combined pools of 90 bumblebee libraries from the same specimens stored using 2 different methods. Bumblebee specimens were sampled between 2017 and 2022 from various locations in Norway, Germany, Denmark and Sweden and were either stored dry at room temperature (n=82) or in ethanol at −18°C (n=8). The box plots compare **(a)** number of reads; **(b)** proportion of collapsed reads; **(c)** endogenous DNA proportion (unique reads only); **(d)** read length and **(e**) coverage.

Finally, when combining the sequencing data of Pool SP and Pool S4, we obtain at least 3-fold overall coverage for 88 out of 90 samples, and at least 10-fold overall coverage for 58. (Table 2).

## Discussion

We here use an affordable, practical and efficient tagmentation-based genomic library preparation protocol to generate successful libraries from 6ng DNA extracted from entomological samples. We reduce costs by 1) transposase enzyme dilution (similar to Jones et al, 2023) and 2) pooling post-amplification libraries without quantification prior to clean-up. We also improve sequencing efficiency by applying a double-sided post-amplification clean-up protocol, selecting a narrow range of library fragments from the broad distribution of fragment sizes produced by the tagmentation reaction. We thus generated medium coverage (>3-fold) genomes for 88 out of 90 specimens, with an average of approximately 10-fold coverage. We estimate the overall economic cost for library preparation and PCR amplification of 96 reactions to be below 450 Euros. Below, we discuss a number of practical considerations.

First, initial extractions of bumblebee samples stored either dry or in ethanol revealed that dry-stored samples produced significantly higher overall DNA yields (up to 3-fold higher, see Figure 1); in fact, few extracts of ethanol-stored samples exhibited DNA concentrations beyond 1ng/μL. Assuming the dry-stored samples to be of higher quality, we chose to prioritise these samples for subsequent DNA extraction, library preparation and sequencing, resulting in the unbalanced sample sizes of the two groups (82 dry-stored samples, 8 ethanol-stored samples; see Table 1). Nonetheless, the eight extracts from ethanol-stored samples performed significantly better during sequencing, despite normalising the concentrations of all samples. Ethanol-stored samples obtained significantly more read pairs, which were also less often fully collapsed, indicating larger DNA fragments. They also yielded significantly greater proportions of endogenous DNA than the dry-stored sample sequences. We speculate that the combination of submerging samples in 96% ethanol with a storage temperature of −18°C reduces the presence of microbial or fungal contaminants, improves the proportion of endogenous DNA and decreases fragmentation of the little amount of DNA remaining. Given that our DNA library preparation protocol is of an adequate sensitivity to produce viable libraries from such low DNA amounts, we conclude that 96% ethanol-based storage of entomological specimens is preferable for high-throughput sequencing compared to dry storage, and that DNA yield is not a determining factor of library sequencing quality.

In order to increase affordability of library preparation, a 6X Tn5 transposase dilution was used, along with direct pooling of all amplified libraries prior to clean-up. As a result of reduced enzyme concentration, the tagmentation reaction produced a broad distribution of DNA library fragment sizes, with a large proportion of long fragments (see Figure 2b; Figure 3, Pool SP). The ideal library fragment size distribution for Illumina DNA sequencing peaks tightly around approximately 600 bp (Illumina DNA Prep Reference Guide, 2020); in addition, long fragments have been found to produce significantly more error rates and reduced base qualities when sequenced on Illumina platforms (Tan et al, 2019). In order to narrow library fragment size distribution around the recommended optimal length of 600 bp and thus increase sequencing efficiency, a more stringent double-sided post-amplification clean-up was performed on Pool S4, which removed both very short (<~400 bp) and very large (>~1000 bp) fragments. Consequently, Pool S4 produced significantly fewer collapsed reads, more endogenous reads and thus yielded approximately 1.5-fold more nucleotides per paired read compared to Pool SP. We therefore conclude that this stringent double-sided clean-up leads to a considerable improvement in sequencing efficiency.

To further reduce financial costs, we pooled all libraries after PCR amplification and omitted individual quantification, which likely increased the variation and spread of coverage obtained by our sequences. Nonetheless, when combining the sequencing data of Pool SP and Pool S4, we find that 88 out of 90 samples have coverage of at least 3X, and 58 out of 90 samples with coverage over 10X. The vast majority of our samples are therefore suitable for a range of downstream analytical approaches, in spite of this ‘quick and dirty’ pooling approach.

Finally, we reiterate that 90 successful libraries were generated from a standardised DNA input of 6ng extracted from just one or two bumblebee legs. This low tissue volume minimises morphological damage to biologically valuable entomological specimens, helps to maintains their potential for future taxonomic, ecological and DNA-based research, and as a result reduces the requirement for additional entomological sampling efforts. In addition, library preparation protocols with low DNA input volumes are essential for the genomic sequencing of much smaller organisms than bumblebees, which may yield very little DNA even when entire specimens are used for DNA extraction. We therefore hope that this protocol can be of considerable value for the future of entomological genomics and monitoring.

## Data availability

All raw sequence read data are available under PRJEB65580 at the European Nucleotide Archive (ENA, https://www.ebi.ac.uk).

## Acknowledgements

Sequencing was performed by the Norwegian Sequencing Centre (www.sequencing.uio.no), a national technology platform hosted by the University of Oslo and Oslo University Hospital, supported by the Research Council of Norway and South-Eastern Regional Health Authority.

## Author contributions

B.S. conceived the study with the assistance of E.M. All samples were provided by M.A.K.S. DNA extraction was done by L.C., and library preparation done by B.S and L.C. Sequencing was done by G.D.G. L.C. performed data quantification and analysis with support from S.K. and M.S., and wrote the manuscript with support from B.S.

## Notes

### Competing Interest Statement

The authors have declared no competing interest.

