## Supplementary Material for "A fast and inexpensive plate-based NGS library preparation method for insect genomics"

### Library protocol for 96-well plates using loaded Tn5 tagmentation

Important notes:
**ALWAYS** spin down plates/tubes before opening
**Prepare the PCR mix** before starting the tagmentation reaction, here we use KAPA HIFI
**ALWAYS** Check your solutions/primers/enzymes.
**Check the chemicals** You need the chemicals for the tagmentation protocol (loaded tagmentation enzyme #C01070012, 2X tagmentation buffer #C01019043, Diagenode), sufficient PCR chemicals, normalized DNA extracts in plate, Illumina primers in plate

**Prepare/defrost buffers**

1) *Defrost 2X tagmentation buffer*
2) *Create **500µL** 0.2% SDS dilution (5µL of SDS 20X stock in 495µL nuclease free water, 1:100 dilution). In a strip tube, divide **8 * 60µL** (480 µL) of 0.2 % SDS*
   *(*NOTE! If SDS solution is "crystallized", heat this buffer to 55-65 ºC to resuspend the solution before use)*
3) *In a strip tube, pipet **8 * 72µL** of 2X Tagmentation buffer (use a 20% overestimate)*

***Prepare DNA***

4) *Spin down DNA 96-well plate with **normalized (1ng/µL) DNA extracts**. Using a multichannel pipet **6µL** of DNA extract of each well to a new plate.*

***Prepare tagmentation enzyme dilutions***

5) *Defrost loaded tagmentation enzyme (Diagenode) and dilute this 1:6 to obtain **120µL** of diluted enzyme; Pipet **60µL** 2X tagmentation buffer and add **40µL** of nuclease free water. Then add **20µL** of tagmentation enzyme and mix. Pipet **14.4µL** of diluted enzyme at each **72µL** 2X buffer in the strip tube. Mix and spin the strip down.*

***Dispense tagmentation enzyme dilution, incubate and stop reaction***

6) *Using multichannel, add **6µL** of Tn5 tagmentation buffer and diluted enzyme mix to the DNA in the plate. Mix by pipetting up and down 4X. Seal the plate and spin down.*
7) *Incubate at **55°C** in PCR machine for 7 min, cool down to 4°C. Spin the plate down.*
8) *Using a multichannel, add **3µL** of 0.2 % SDS (from 8 tube strip) to each well. Mix by pipetting 4X, seal and spin down. Leave for 5 minutes at room temperature (stopping tagmentation reaction).*

Tagmentation reaction, 12µL per reaction + 3 µL SDS

| | Volume (µL) |
|---|---|
| DNA (6 ng, 1ng/µL) | 6 |
| Tn5 buffer (2x) | 5 |
| Tn5-oligo complex (diluted 1:6) | 1 |
| After 55ºC, 7 min step: | |
| 0.2% SDS (1:100 from 20%stock) | 3 |

Total reaction volume is 15ul, this is directly used in the *post-tagmentation PCR*

#### Post-tagmentation PCR

Prepare PCR Mastermix (prepare this beforehand and place in fridge)

1) In **two** 1.5 ml tubes: pipet **770µL** (55 * 14) of Nuclease free water
2) Add **444µL** of KAPA HIFI buffer (55 * 8) each of the two tubes
3) Add **66µL** dNTP mix (55 * 1.2) each of the two tubes
4) Add **44µL** KAPA HIFI polymerase (55 * 0.8) to each of the two tubes. Mix well and spin down.
5) Divide **162µL** of Mastermix in 16 slots over two strip tubes (2X 8 slots). Store in fridge.

Dispense PCR Mastermix and add primer combination

1) Using a multichannel, pipet **24µL** of Mastermix into each tagmentation DNA slot. Mix by pipetting up and down 2 times.
2) Make SURE to spin the primer plate down before opening. Using a multichannel, add **1µL** of N7 and N5 primer to each well. Mix by pipetting up and down 5 times. Carefully tape the plate. Make sure you know the orientation of the primers (A1 should go onto A1 sample). Spin the plate down and place in PCR machine.

##### post-tagmentation PCR reaction

| | Per reaction | X |
|---|---|---|
| Tagmentation DNA | 15 | |
| Nuclease free H20 | 14 | |
| **KAPA HiFi buffer** | 8 | |
| **dNTP mix, 10mM** | 1.2 | |
| **KAPA HiFi polymerase** | 0.8 | |
| N7and N5 primer 1uM | 1 | |
| Total volume | 40 | |

72°C for 3min

95°C for 3min

**12x** (98°C for 20sec, 55°C for 30sec, 72°C for 30sec.)

##### Pool the plate PCRs:

1. After the PCR reaction, spin the plate down, and using a multichannel pipet, combine 6µL of each well ***per row*** to an 8-strip tube. (i.e., 12 columns are combined into a single tube) Mix the strip well and spin down.
2. Take 65µL of each 8 tube and pipet into a single Eppendorf tube. Mix well. Divide 120µL into 4 tubes (of an 8-strip tube). Proceed to clean-up.

#### AMPure XP purification (2x!)

*Use a fairly aggressive clean-up to get to the right size distribution of the library pool. Do a **right** (large fragments) and **left** (small fragments) side profile cleanup, TWICE. This costs a LOT of DNA, so we concentrate the libraries over 10-fold during the cleanup*

***First** clean-up, 4 tubes*

1. Add 60 µL of Ampure XP (0.5 ratio large fragments cleanup) to the 120µL reaction in each tube. Incubate for 5 minutes at room temperature.
2. Set the 4-tube strip on the magnetic plate and wait 1-2 minutes for the beads to collect on the side.
3. NOTE: **Keep** the supernatant, pipet 180 µL to NEW tubes. Throw away the tubes with beads. Add **18µL** of Ampure beads to the supernatant (0.65 small fragments cleanup).
4. Set the 4-tube strip on the magnetic plate and wait 1-2 minutes for the beads to collect on the side. Remove the supernatant.
5. Add 200µL 80% ETOH (make fresh!) and wash the beads. Keep the beads on the plate, 30 second incubation is enough, so just proceed.
6. Remove the ETOH and repeat.
7. After removal of last ETOH, let the samples stand ~5-8 min in order for the beads to dry (but not too much!)
8. Elute in 52µL H2O. Let stand for 2minutes at room temperature and then put on the magnetic block in order to remove supernatant containing DNA to a new tube. Combine two tubes into one to obtain **100µL** DNA extract.

***Second** clean-up, 2 tubes*

1. Add **50µL** of Ampure XP (0.5 ratio large fragments cleanup) to the 100µL reaction in each tube. Incubate for 5 minutes at room temperature.
2. Set the 2-tube strip on the magnetic plate and wait 1-2 minutes for the beads to collect on the side.
3. NOTE: We **keep** the supernatant, pipet 150 µL to NEW tubes. Throw away the tubes with beads. Add **15µL** of Ampure beads to the supernatant (0.65 small fragments cleanup).
4. Set the 2-tube strip on the magnetic plate and wait 1-2 minutes for the beads to collect on the side. Remove the supernatant.
5. Add 200µL 80% ETOH and wash the beads. Keep the beads on the plate, 30 second incubation is enough, so just proceed.
6. Remove the ETOH and repeat.
7. After removal of last ETOH, let the samples stand ~5-8 min in order for the beads to dry (but not too much!)
8. Elute in **20µL H2O**. Let stand for 2minutes at room temperature and then put on the magnetic block in order to remove supernatant containing DNA to a new tube. Combine both tubes into a single **~40µL** pool This is your pool for sequencing. Label the pool well.

**Check the library on the Fragment Analyser.**

Example of a good size distribution:

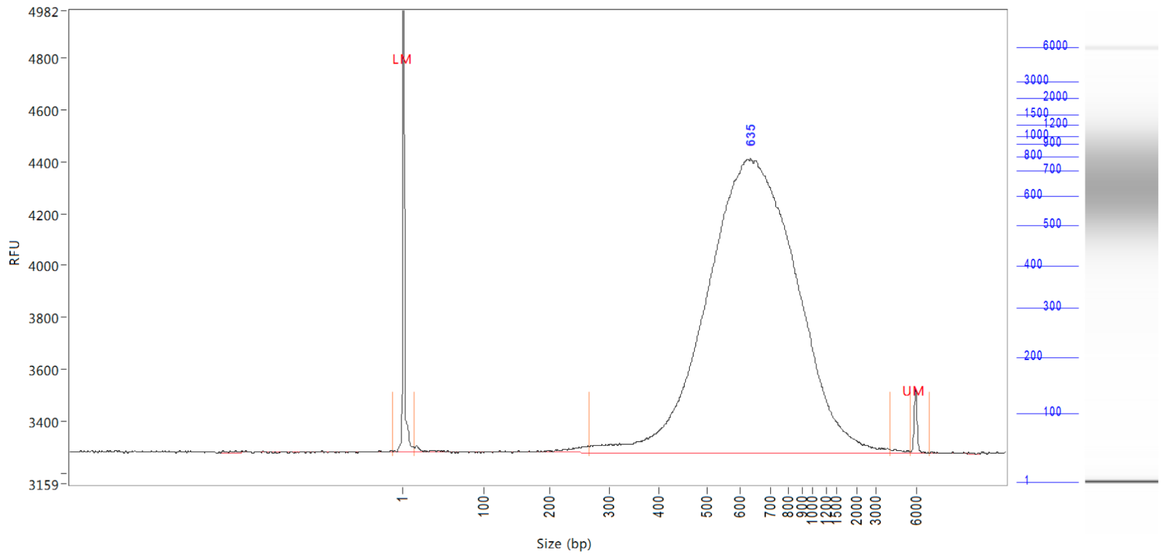

**KEEP the uncleaned libraries frozen and safely stored, well-labeled. If planned well and ahead, the protocol (and FA) can be run in a single day.**
